# Mechano-activated chondroprogenitors (MACs) drive intrinsic cartilage repair; a process that is arrested in osteoarthritis

**DOI:** 10.1101/2025.08.21.671507

**Authors:** Linyi Zhu, Hayat Muhammad, Thomas A Perry, Anastasia Ardasheva, Rebecca A Symons, Jessica J McClure, Karolina Kania, Susan M Clark, Fabio Colella, Lada A Koneva, Sumayya Khan, Suzanne Elizabeth Eldridge, Francesco Dell’Accio, Simon Mastbergen, Mylène Jansen, Fiona E Watt, Yoshi Itoh, Jane Bennett, Jadwiga Zarebska-Miotla, Moustafa Attar, Chris Chan, STEpUP OA Consortium, Anke J Roelofs, Cosimo De Bari, Stephen N Sansom, Jana Riegger, Tonia L Vincent

## Abstract

Articular cartilage, an avascular, matrix-rich tissue, is thought to have limited repair, thereby contributing to osteoarthritis (OA), the common degenerative disease of joints. Cartilage regeneration does occur, however, in OA joints that are mechanically off-loaded. Here we show that mechanical stress, through release of matrix-bound growth factors, reprogrammes chondrocytes, the primary cells of cartilage, into ‘mechano-activated chondroprogenitors’ (MACs). Studying OA joint fluid before and after mechanical off-loading, reveals evidence of chronic MAC activity, which switches back to a chondrogenic one when mechanical stress is removed. Taken together, we conclude that OA is a disease of ‘arrested repair’ in which mechanical stress signals need to be switched off before full repair can occur. This novel paradigm uncovers exciting new treatment opportunities.

## INTRODUCTION

Over a third of adults will develop osteoarthritis (OA), a disease with huge societal burden and for which no disease modifying treatments exist (*1-3*). The disease is characterised by breakdown of the articular cartilage, a matrix-rich, avascular tissue, which supports chondrocytes, the primary cell of cartilage. Mechanical stress, either through micro- or macro-injuries, is a common aetiological factor in the development of OA (*4, 5*), with chondrocytes displaying exquisite mechanosensitivity to injurious and physiological mechanical loads (*6*). One important mechanism of mechanotransduction is through a rapid (<5 min) sodium-dependent displacement of growth factors from the heparan sulfate binding sites of perlecan, located in the pericellular matrix (PCM) of cartilage (*7*). Several injury-released growth factors have been identified by our group using mass spectrometry, including fibroblast growth factor-2 (FGF2), connective tissue growth factor (CTGF), covalently bound to latent transforming growth factor beta (TGFβ), hepatoma derived growth factor (HDGF) and Cysteine-rich angiogenic inducer 61(CYR61) (*7-9*). The chondroprotective role of FGF2 and CTGF-TGFβ has been demonstrated in vivo (*10, 11*). HDGF and Cyr61 have reported pro-regenerative activities in other tissues (*12-14*).

Poor repair capacity in cartilage has been attributed to the paucicellular and avascular nature of the tissue, although cartilage regeneration has been observed in individuals with advanced OA who have undergone surgical joint distraction (SJD); a procedure in which the joint is mechanically off-loaded using an external fixator for a period of 6 weeks (*15-18*). It has also been observed in mice following focal cartilage injury in an age and strain-dependent manner (*19*). Putative repair cells have been expanded from healthy and osteoarthritic human cartilage (*20-24*) raising the possibility that they originate from within cartilage, driving so-called ‘intrinsic repair’. Here we show that primary chondrocytes give rise to intrinsic repair cells of cartilage and investigate how they are activated and why chronic activation needs to be switched off before full repair can occur.

## RESULTS

### Chondrocytes contribute to cartilage repair

We first tested the hypothesis that articular chondrocytes contribute directly to cartilage repair after focal cartilage injury in vivo, using young mice in which Col2-lineage cells (chondrocytes) had been labelled at 2 weeks of age using a Cre^ER^-inducible Td tomato transgene (Fig. 1A, 1B). Eight weeks after cartilage injury, Col2-lineage cells were found in repair tissue at a percentage similar to the adjacent intact cartilage (47.5 ± 13.7, 58.3 ± 6.9 respectively) (Fig. 1C, 1G), whilst being less abundant in subchondral bone marrow (Fig. 1D) and synovium (Fig. 1E). These data suggested that articular chondrocytes are a major source of repair cells after injury, driving intrinsic cartilage repair.

**Fig. 1.**
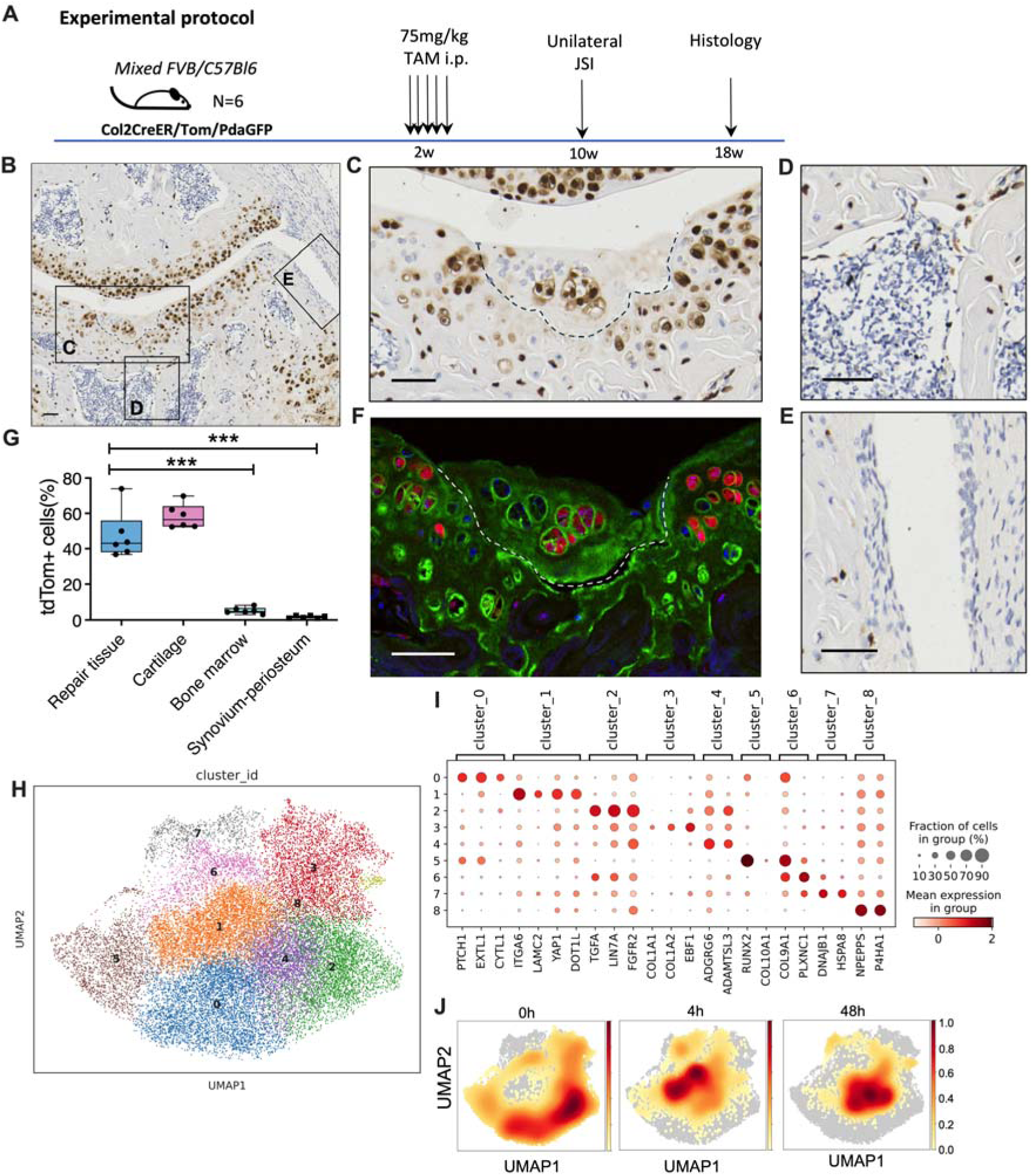
Intrinsic repair cells of cartilage upon injury. **A**, Schematic showing the protocol for Col2 lineage cell tracing in female *Col2-CreER;tdTom* mice undergoing joint surface injury (JSI). **B**, Representative image of *Col2*-lineage cells in knee joint of *Col2-CreER;tdTom* mouse 8 weeks after focal cartilage injury detected by tdTom immunohistochemistry with **C**, region of defect, indicated by dotted line (also enlarged), **D**, subchondral bone marrow and **E**, synovium and periosteum shown. **F**, Defect region also shown with immunofluorescence signal for tdTom (red) and type II collagen (green). Scale bars = 50 µm. **G**, Quantification of % of tdTom-expressing *Col2*-lineage cells in each joint tissue (n = 6). Mean ± 95% confidence interval. *** p<0.001. (H-J) Single-nuclei RNA-seq of porcine articular cartilage after injury. **H**, Uniform manifold approximation and projection (UMAP) plot of all chondrocytes (n = 25,075 nuclei) coloured by cluster. **I**, Dot plot of the top positive marker genes for each cell cluster with % of cells expressing marker gene and average expression level shown. **J**, Density of cells across clusters at each time point (0h, n=4; 4h, n=3; and 48h, n=3 cartilage samples). For gene identities see data file S1.

To understand how the chondrocyte gives rise to repair cells after injury, we developed a method for single nuclei RNA-sequencing (snRNA-Seq) from healthy porcine cartilage before (time 0) and after mechanical cutting injury (4 and 48h). The cluster structure of all the samples is presented for the whole-tissue analysis, including cells of the osteochondral junction (fig. S1A). The majority (81.9%) of nuclei analysed were deemed to be from chondrocytes, with the remaining cells identified as endothelial (4.0%), bone (3.7%), neuronal-like (8.2%) and monocyte/macrophage (2.2%), by marker gene profiles (fig. S1B, S1C). Their cluster composition changed over time (fig. S1E). The cluster structure of chondrocytes alone and their associated marker genes are shown in Fig. 1H, I. Of the nine chondrocyte clusters defined, most cells (85.5%) were in clusters 0, 2, 3, and 5 prior to injury (time 0) (Fig. 1J, fig. S2A). At 4h after injury, chondrocytes assumed a new phenotype, cluster 1 (49.0%), which was associated with reduced cell numbers in clusters 0, 2, and 5. By 48h there was a partial shift into cluster 4 (Fig. 1J, fig. S2A). The marker genes identified for cluster 1 included integrin alpha 6 (*ITGA6*), laminin C2 (*LAMC2*), Yes-associated protein-1 (*YAP1*), and DOT1-like histone lysine methyltransferase (*DOT1L)*, which were all highly expressed in this cluster (Fig. 1I). Cluster 1 was also marked by low expression of vitrin (*VIT*), TGFβ receptor 3 (*TGFBR3*), and cartilage oligomatrix protein (*COMP*) (data file S1). Cluster 1 genes were regulated by cartilage injury (at 4h) when considering the pseudobulk level analysis of all chondrocytes (fig. S2B). In total, we identified 1,350 up-regulated genes and down-regulated 1,256 genes at 4h (DESeq2 analysis, Benjamini-Hochberg (BH) corrected p <0.05, |Fold change| >1.5) (data file S2).

### PCM growth factors drive chondrocyte epithelial mesenchymal transition (EMT)

We sought to determine whether the early phenotypic switch in chondrocytes following cartilage injury was driven by release of PCM-bound growth factors. Accordingly, medium conditioned for 5 min by cutting porcine cartilage (injury CM), containing PCM-derived growth factors (*7, 9*), was used to stimulate isolated primary porcine chondrocytes. Positive cluster 1 marker genes were up-regulated in response to injury CM and this was dependent on FGFR signalling (Fig. 2A). Bulk mRNA sequencing identified 4,007 genes that were significantly regulated by injury CM (DESeq2, |Fold change| >1.5, BH adj p<0.05) (Fig. 2B, data file S3), of which 846 were significantly changed with FGFRi (fig. S3A, S3B, data file S4). The genes regulated by injury in the pseudobulk analysis of the snRNAseq data (data file S2) positively correlated with genes regulated in chondrocytes by injury CM (data file S3) as shown in fig. S3C (r=0.406, p<2.2 10E-16), but negatively correlated with injury CM plus FGFRi (data file S4) in fig. S3D (r=-0.322, p<2.2 10E-16). Genes regulated by injury CM were enriched for Hallmark pathways associated with inflammatory signalling (“TNF signalling”, “inflammatory response”) and “Epithelial to Mesenchymal Transition (EMT)” (FGSEA analysis, BH padj <0.05) (Fig. 2C, table S1), in an FGFR-dependent manner (Fig. 2D, table S2). EMT pathway genes included some of the most strongly FGFR-dependent genes after injury CM stimulation, including *LAMC2* and *INHBA*, the dimer of which forms activin A, a TGFβ family member (Fig. 2B).

**Fig. 2.**
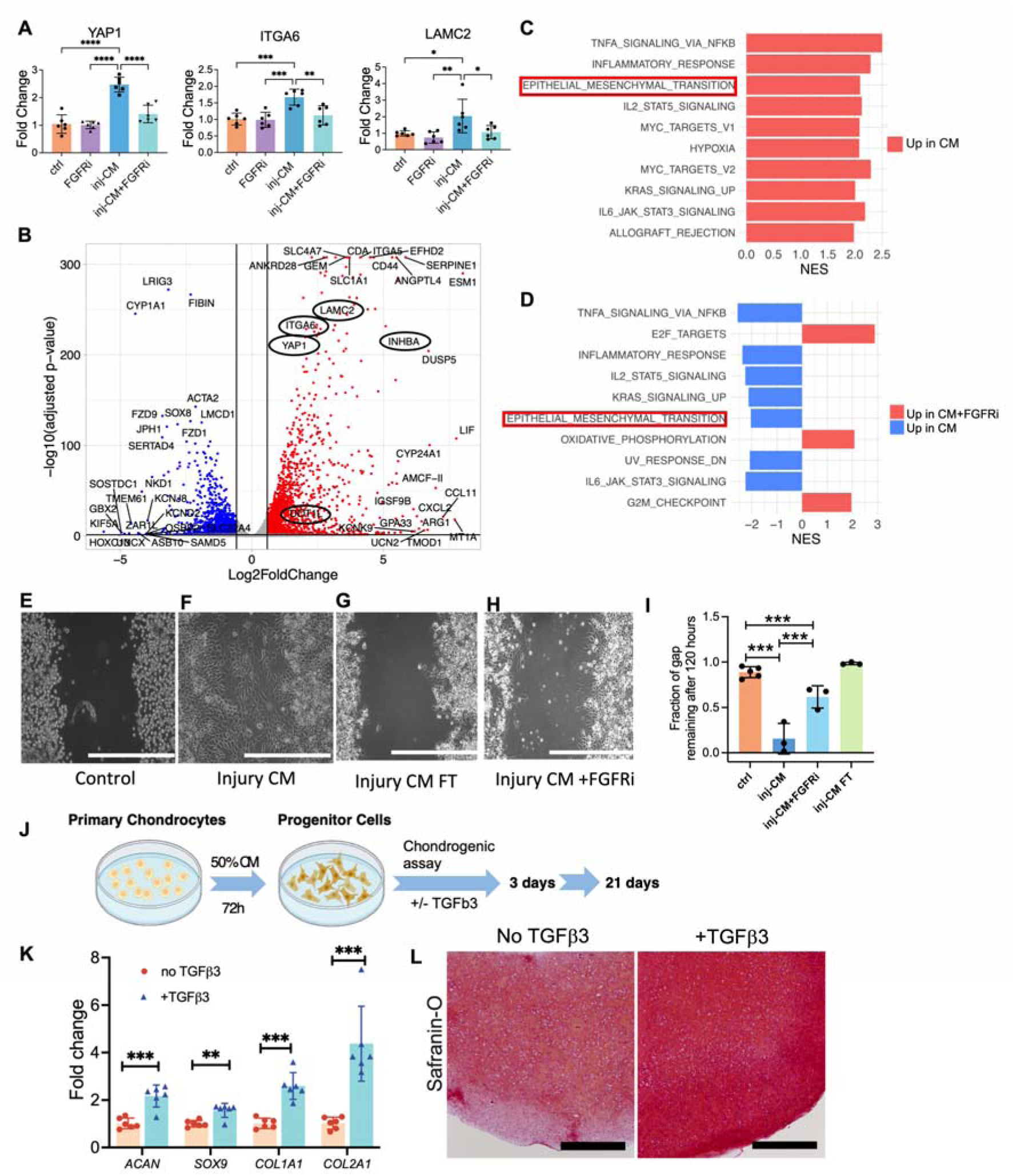
Injury CM drives epithelial to mesenchymal transition (EMT) to reprogramme chondrocytes into MACs. **A**, qPCR was performed on primary porcine chondrocytes after stimulation with injury CM with and without FGFRi, n = 6. **B,** Volcano plot from bulk RNA-seq analysis showing differentially expressed genes in injury CM stimulated vs unstimulated chondrocytes (Deseq2, padj < 0.05, |fold change| > 1.5). Top genes are labelled. **C,** Top 10 pathways by gene enrichment analysis (Hallmark) of injury CM vs ctrl and **D,** injury CM with and without FGFRi (padj < 0.05). **E-H** Representative images of the chondrocyte ‘gap assay’ after different stimulations (120h) including injury CM flow through (FT) from heparin column (Injury CM FT), scale bar = 500 µm. **I**, Quantification of images in **E-H**, n=3-5. **J**, Primary chondrocytes stimulated with injury CM for 72 h then stimulated with defined chondrogenic medium with (+TGFβ3) or without (no) TGFβ3. **K**, The expression of chondrogenic marker genes at day 3. **L**, Safranin-O staining of the chondrogenic pellets at day 10, scale bar = 200 µm. One-way ANOVA with Bonferroni post-hoc analysis. Bars represent the means ± SEM. \**P* < 0.05, \*\**P* < 0.01, and \*\*\**P* < 0.001.

EMT is the cellular process by which cells leave their matrix attachments to become motile and proliferative (*24*). It is implicated in development, tissue repair and cancer and has been linked previously to YAP1, ITGA6, LAMC2, DOT1L and INHBA (*25-31*). Treatment of confluent primary chondrocytes in a ‘gap’ assay with injury CM led to chondrocyte elongation and migration, leading to gap closure (Fig. 2E, F). This was driven by heparin binding factors, as passing the injury CM through a heparin column removed activity and was partially FGFR-dependent (FGFRi) (Fig. 2G-I). The response was also captured and quantified by live imaging after stimulation of cells at low density with injury CM (Fig. 3A, B) (Video data file S5). Neutralising antibodies to FGF2 inhibited cell density to a similar degree as FGFRi, confirming that FGFR signalling is driven by FGF2 (Fig. 3C). The importance of FGF2 was also demonstrated in a YAP-TEAD reporter cell line (*32*), where activation of Yap with injury CM was also FGFR-dependent (Fig. 3D). Furthermore, reduced cartilage repair was observed in vivo in FGF2^-/-^ mice (Fig. 3E, F).

**Fig 3.**
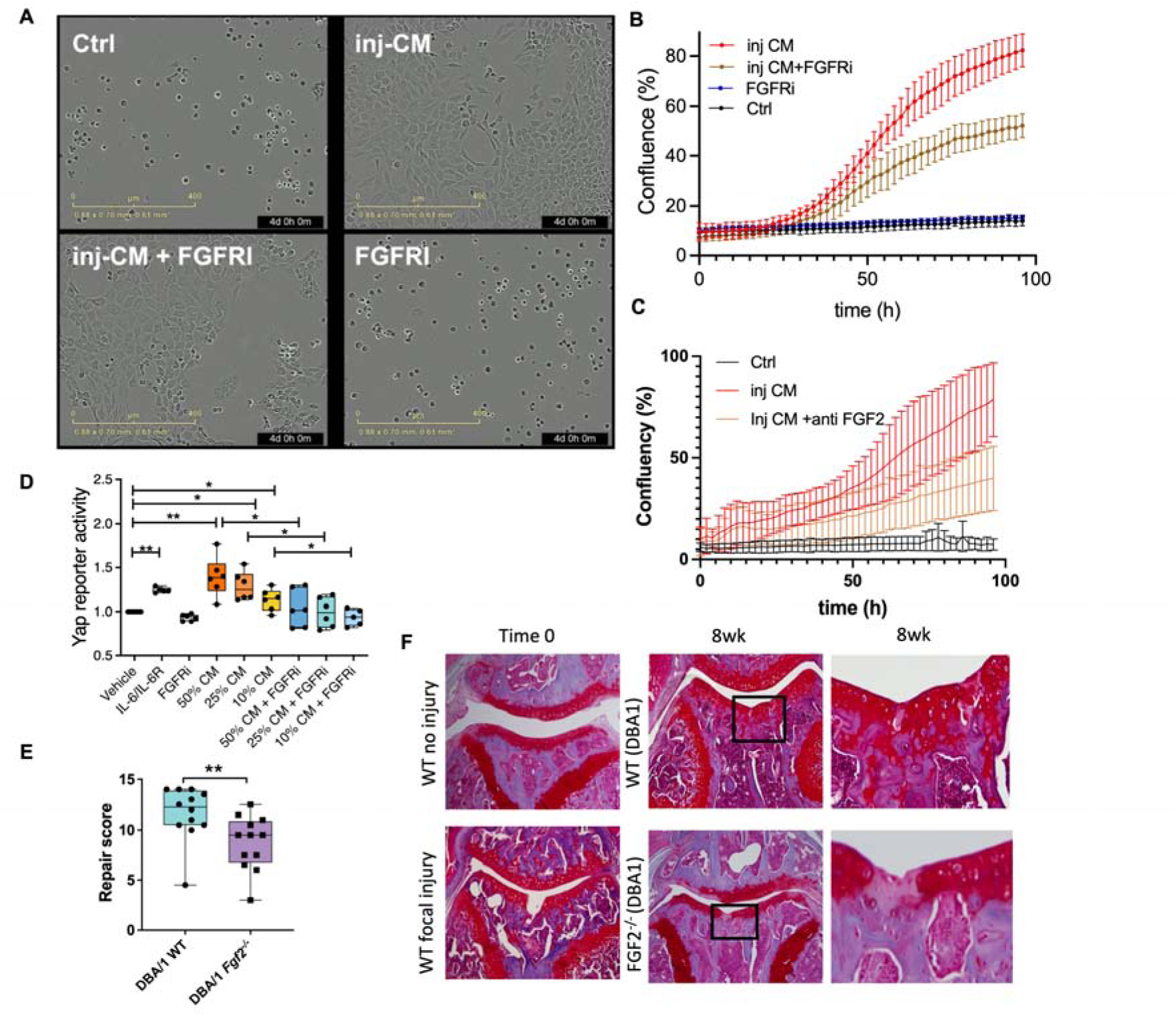
FGF2 contributes to MAC generation, YAP1 activity and in vivo cartilage repair. Low density porcine chondrocytes were stimulated with injury CM in the presence or absence of FGFRi (250nM SB402451) or anti-FGF2 neutralising antibody (4 μg/mL) and imaged using the IncuCyte imaging system. **A**, Representative images of cells after 96h culture. **B**,**C**, Quantitation of cell density assessed by IncuCyte imaging (at 2h intervals) over 96h. Movie showing change in cell morphology and proliferation for each condition are presented in supplementary video data file S5. **D**, Yap-Tead GFP reporter assay after stimulation of cells with IL-6/IL-6R (positive control), injury CM at indicated dilutions, with or without FGFRi (250 nM). Graph shows mean ± SD. Data points indicate independent experiments. One-way ANOVA with Tukey’s post-test, n = 6. Quantification **E**, and representative images **F**, of in vivo murine cartilage repair model, induced at 10 weeks of age and assessed after 8 weeks, in FGF2^-/-^ and DBA1 control mice. Histological sections were scored using a modified Pineda score. Mean ± SEM; Mann-Whitney U test; *p>0.05, **p>0.01, n = 12 biological replicates.

Using a chondrogenesis pellet assay in which cells are stimulated with defined chondrogenic medium containing TGFβ3 (*33*), chondrocytes activated by injury CM (Fig. 2J) were able to return to fully differentiated (homeostatic) chondrocytes as shown by increased chondrogenic gene expression (Fig. 2K) and characteristic histological appearance (Fig. 2L). Collectively, these data reveal that PCM-derived growth factors, released upon injury, lead to ‘mechano-activated chondroprogenitors’ (MACs) with intrinsic cartilage repair potential.

### MACs drive late cartilage repair response

In native cartilage, the repair benefit of MAC generation may not be realised unless the cells can be switched back to homeostatic chondrocytes. We hypothesised that a single acute injury would lead to spontaneous reversion of the MAC to chondrocyte once the acute injury response had resolved (soluble growth factors bind back into the PCM). We employed a cartilage drop tower model in which injured or uninjured cartilage was cultured for 2 weeks following a single high-impact injury (Fig. 4A, B). Like cutting injury, impact injury led to a transient FGFR-dependent increase in *YAP1, LAMC2, ITGA6* and *INHBA,* 4h after injury in the bulk tissue (Fig. 4C). New collagen synthesis was first observed 7 days after injury (*COL1A1*) which shifted to collagen type II a1 chain *(COL2A1*) at day 14 (Fig. 4C), indicative of chondrogenesis. Both collagen responses were inhibited by FGFRi indicating that the early acute injury leads to a subsequent late chondrogenic (anabolic) response in the injured tissue. By performing snRNA-Seq of the tissue 14 days after injury, discernible chondrocyte clusters were once again revealed (Fig. 4D, fig. S4), but now no distinct injury clusters were defined (Fig. 4E). Gene set enrichment analysis of genes differentially expressed in response to injury (D14) (pseudobulk analysis DESeq2, BH padj <0.05 and |Fold change| >1.5) revealed a strong anabolic (chondrogenic) signature (Fig. 4F, highlighted), with select differentially expressed genes labelled in Fig. 4G (data file S6).

**Fig 4.**
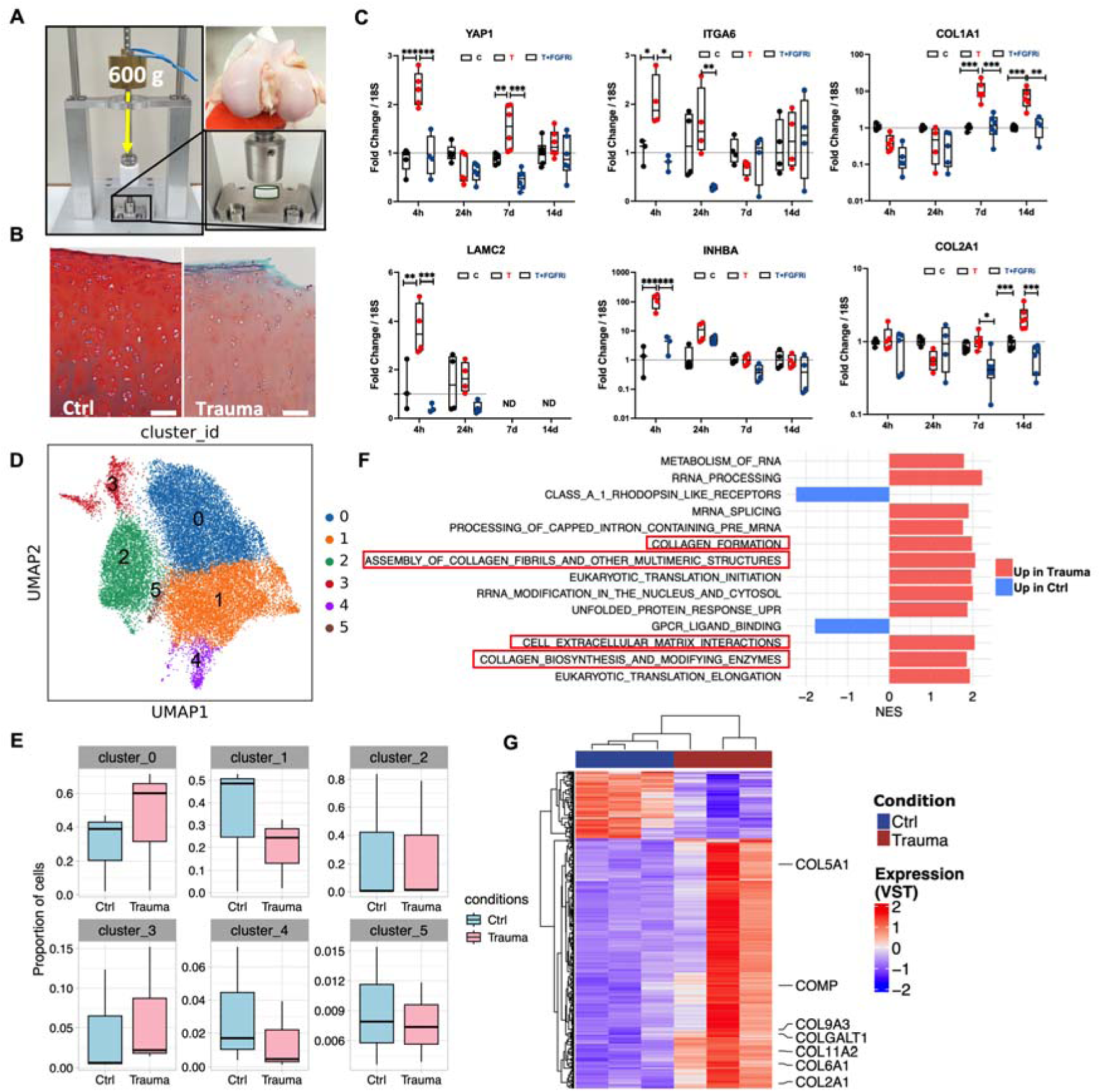
Acute impact injury to cartilage leads to a late (14 day) anabolic tissue response. **A**, Porcine knee articular cartilage explants were dissected from the joint, rested for 48h then either subjected to a single high-impact load (Trauma) by a drop tower, or were left uninjured (Control, Ctrl), in the presence or absence of FGFRi. **B**, Histological assessment of cartilage by Safranin-O staining after 14 days. **C**, qPCR was performed on bulk tissues over the 14-day period and analysed for *YAP1, ITGA6, LAMC2, INHBA, COL1A1 and COL2A1.* C: control, T: Trauma. **D**, snRNA-Seq of porcine articular cartilage 14 days after injury. The UMAP plot for all chondrocytes (*n* = 24,022 nuclei), coloured by cluster is shown. **E**, Proportion of cells in each cluster was calculated as percentage of the total cells at each condition. **F,** Gene enrichment analysis (Reactome) of differentially expressed genes using the pseudobulk counts (FGSEA analysis, BH adjust p < 0.05). **G**, Pseudobulk heatmap showing examples of cartilage anabolic genes regulated in the tissue after injury (Deseq2 analysis, BH adjust p < 0.05, |fold change| > 1.5).

### Chronic MAC activity suppresses repair

OA is known to be a disease of chronic mechanical stress (*4*), which could result in uncontrolled MAC activity with suppression of chondrogenesis. Indeed, this proved consistent with a recently completed large-scale proteomic analysis in 1,361 synovial fluid samples from individuals with knee OA, which identified “EMT” as the dominant pathway associated with severity of OA; an activity that was robust even after stratification by age, body mass index, or biological sex (*34*) (in revision). Of note, the lead proteins driving EMT in synovial fluid were inhibin bA, its homodimer, activin A, and its heterodimer activin AC, which were also among the top 10 proteins most strongly associated with radiographic disease severity (*34*). We hypothesised that mechanical off-loading of the joint would result in switching off injury-induced mechanical signalling and allow chronically activated MACs to rebuild their matrix and regain their homeostatic chondrocyte phenotype. Mechanical joint off-loading can be achieved through ‘surgical joint distraction’ (SJD) in which the joint is ‘pulled apart’ by an external metal frame secured by pins on either side of the joint (Fig. 5A). Individuals mobilise with the device in situ for 6 weeks, then it is removed, and they are rehabilitated. SJD leads to sustained improvement in pain out to 2 years following removal of device with evidence of cartilage regrowth on magnetic resonance imaging(*16, 35*). To test our hypothesis, we performed proteomic analysis, using the SomaScan platform, of 16 paired synovial fluid samples taken from individuals with established OA, before and at the end of 6 weeks of surgical joint distraction. 1,576 proteins were significantly changed by joint distraction (Fig. 5B). Inhibin βA, activin A, and activin AC were amongst the strongest down-regulated proteins, along with other TGFβ family members including bone morphogenetic proteins 5 (BMP-5) and 6 (BMP-6) (Fig. 5C, upper panel, data file S7). Joint distraction also led to up-regulation of several proteins associated with extracellular matrix organisation (Fig. 5D) including key chondrogenic marker proteins, SRY box transcription factor (SOX9), COL2A1 (just failing to reach padj<0.05), COL10A1, HPLN1 (hyaluronan and proteoglycan link protein 1, also known as link protein) as well as the PCM protein, HSPG2 (heparan sulfate proteoglycan-2, also known as perlecan) (Fig. 5C, lower panel). Finally, we examined the specific role of activin A in this process. Neutralising antibodies targeting activin A did not alter chondrocyte behaviour following stimulation of primary chondrocytes with injury CM (Fig. 5E), but activin A strongly suppressed chondrogenesis in MACs when cultured with defined chondrogenic medium containing TGFβ3 (Fig. 5F); a behaviour that had previously been described for FGF2 in mesenchymal stem cells (*36*). Activin A therefore appears to be important in maintaining ‘stemness’ of the MAC pool and suppressing their re-differentiation into chondrocytes.

**Fig. 5.**
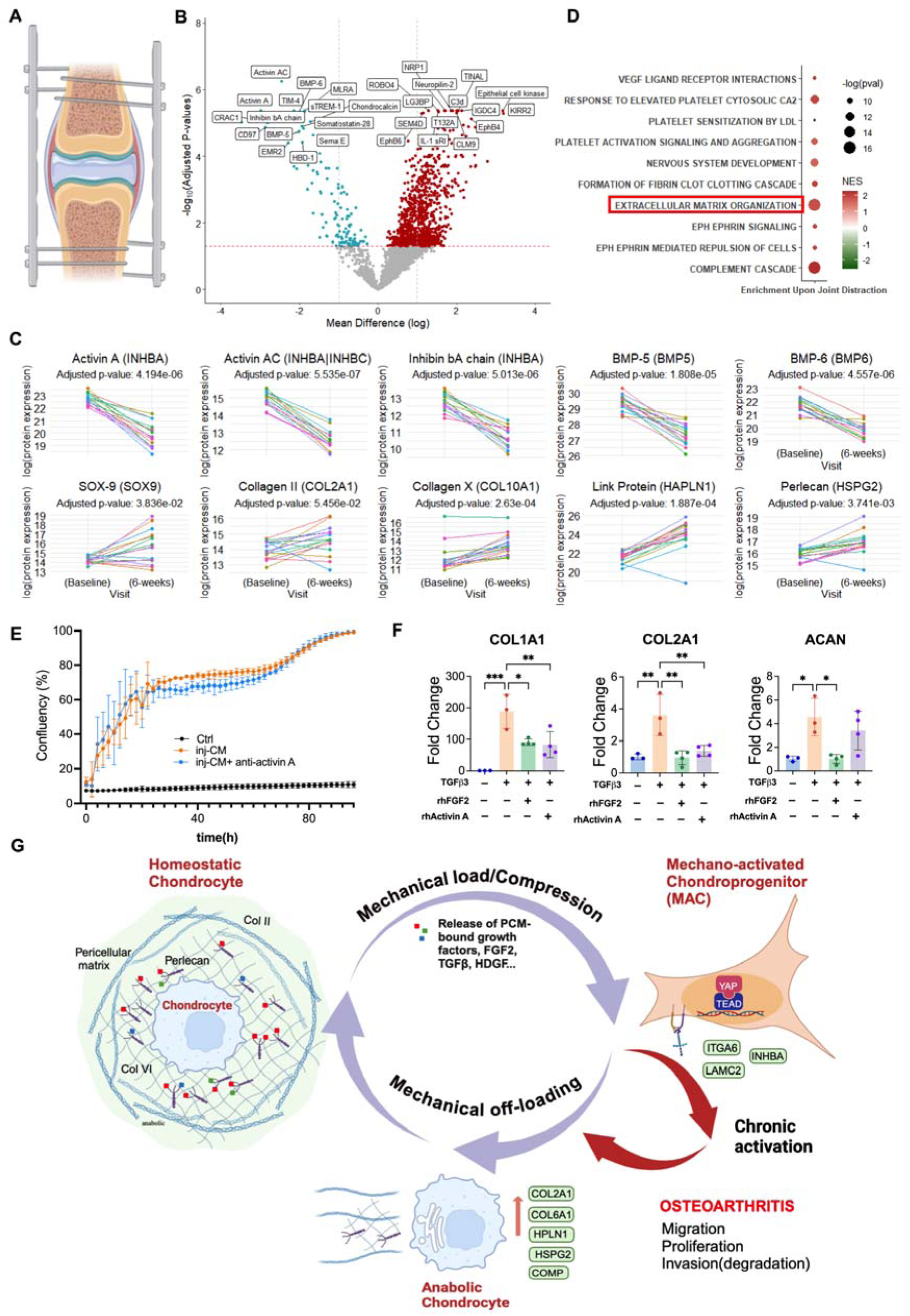
Mechanical off-loading of the knee joint by surgical joint distraction (SJD) changes the synovial fluid proteome and identifies activin A as a suppressor of chondrogenesis in OA. **A**, Illustration of SJD. **B,** Volcano plot showing mean difference (log) in protein expression against adjusted p-values, pre and post SJD in paired samples (n = 16, 5471 proteins, padj ≤ 0.05). Top regulated proteins labelled. **C,** Line plots for a selection of regulated proteins. INHBA, inhibin βA; ACTIVIN A, inhibin βA homodimer; ACTIVIN AC, Inhibin βA:Inhibin βC heterodimer; BMP, bone morphogenetic proteins-5 & 6. SOX-9, SRY box transcription factor-9; COL2A1, collagen type II a1 chain; COL10A1, collagen type 10 a1 chain; HPLN1, link protein; HSPG2, perlecan. **D**, Top 10 pathways after gene enrichment analysis (Reactome). **E**, Cell confluency by IncuCyte imaging of primary chondrocytes stimulated with injury CM in the presence of anti-activin A. **F**, Chondrogenesis assay of MACs (generated by 72h treatment with injury CM) in the presence of recombinant FGF2 or activin A. Expression of chondrogenic marker genes measured by qPCR at day 3. **G**, Schematic of the intrinsic repair cycle of articular cartilage. A and G created using BioRender.

Taken together, these data strongly support a new paradigm in intrinsic cartilage repair that requires two critical components: the first involving growth factor-mediated reprogramming of chondrocytes into motile and proliferative MACs, and the second requiring MACs to re-differentiate back into chondrocytes, rebuild their local matrix (anabolic chondrocyte) and restore homeostasis, a process that is suppressed in OA by chronic growth factor-dependent injury signals (Fig. 5G).

### Discussion

The question of whether cartilage has capacity to repair has been long debated. Historically, two types of articular cartilage repair have been proposed; one that was derived from cells of the cartilage (intrinsic repair) and one that was derived from cells that migrated into cartilage from other tissues (extrinsic repair) (reviewed in (*37*)). Intrinsic repair was thought to produce higher quality tissue but was less commonly observed, especially in the presence of a florid extrinsic cell response (*38, 39*). The data that we present in this paper align with the intrinsic repair that was observed over 60 years ago. Here, we show that repair cells derive from the primary chondrocyte; we define the molecules that initiate their reprogramming and we recognise the need for these pathways to be switched off in order for the cartilage repair cycle to complete.

FGF2 is identified as a key mediator of the repair response, albeit acting in concert with other injury-released growth factors. FGF2 appears to have three distinct roles in this process: it is a key driver of EMT, including YAP activation, previously shown to be important in cartilage repair (*40*); it induces important genes such as *INHBA* and *DOT1L,* the latter which protects against OA and is an OA GWAS hit (*41*); and it suppresses re-differentiation of the MAC back into a chondrocyte (either directly or via activins). Together these maintain the repair cell pool until conditions are favourable for them to re-differentiate into chondrocytes. This accords well with the seemingly paradoxical chondroprotective and anti-chondrogenic actions of FGF2 that have been reported (*11, 42-44*). It also supports why intermittent dosing of sprifermin, a truncated form of FGF18, increases cartilage thickness in patients with OA (*3*), while continuous stimulation with FGF18 supressed chondrogenesis (*23, 45*). The role of the other growth factors in the injury response is less clear. CTGF, covalently bound to latent TGFβ, controls the bioavailability of TGFβ, but only when the target cell expresses TGFBR3 (*8*). Levels of TGFBR3 are suppressed in MACs (cluster 1), suggesting that TGFβ activation is prevented, as might be expected if chondrocyte differentiation is to be delayed. Active TGFβ is likely to be a key driver of chondrogenic differentiation in repairing cartilage (*46*).

Importantly, in OA, our data suggest that continued mechanical loading of the damaged joint leads to chronic activation of MACs and suppression of chondrogenesis. This will lead to further damage as MACs proliferate, migrate and invade (degrade) the surrounding matrix. The biological response to joint distraction strongly suggests that it is possible to switch off chronic injury signals and trigger chondrogenesis and cartilage repair. This may in part be mediated by a drop in activin A. It may also be driven by enhanced signalling through cell guidance pathways e.g. ephrins (Fig. 5D), which have diverse functions including inhibition of mobilisation, promotion of cell differentiation and cellular repulsion (*47-49*). The re-differentiating chondrocyte must now reassemble its PCM (perlecan) and build back the extracellular matrix.

Taken together our data uncover the remarkable ability of cartilage, an avascular tissue, to repair itself by dynamic reprogramming of the chondrocyte in response to sequestered growth factors released by external mechanical stimulation. Repair in OA is observed when the joint’s mechanical environment is directly improved, but it is possible that other more accessible physical or pharmacological therapies could also be exploited to benefit patients.

## MATERIALS AND METHODS

### Study design

This study investigates the role of mechanical stress in chondrocyte reprogramming and OA using healthy porcine and mouse articular cartilage alongside human OA synovial fluid before and after mechanical joint off-loading. Sample size for the chondrocyte and cartilage gene expression experiments were based on previous results from our team where 3 to 6 samples per group are sufficient to show robust regulation of genes upon injury. Joint distraction numbers (n = 20) were previously validated (published) for a small set of proteins by immunoassay showing significant changes in proteins upon treatment. Numbers for snSeq of injured cartilage were a priori set at n = 3-4 assuming that strongly regulated injury response genes would show significant regulation. All sample sizes are annotated within the respective figure legends. For SomaLogic protein analysis, only participants with paired SF samples at baseline and 6 weeks were included. A single pair was excluded due to issues with protein reads; leaving 16 pairs (n = 32 samples). Synovial fluid samples were randomised to plate for molecular analysis by SomaLogic. In vivo repair studies included n = 6 (Col2-lineage) and n = 12 (FGF2^-/-^) mice, based on published work. All mice underwent the same focal cartilage injury surgery. Genotypes were separated by cage. All in vivo data were analysed by two independent blinded scorers. Male (FGF2^-/-^ study) and female mice (Col2 lineage study) were used; prior published studies confirming no sex-dependent repair differences.

### Reagents and Antibodies

Human recombinant FGF2 was purchased from Peprotech EC Ltd. Recombinant human Activin A (MAB3381), TGF-βR3 (QK054), FGF2 neutralizing antibody (AF-233-NA) and Activin A neutralizing antibody (MAB3381) were obtained from R&D Systems. The FGF receptor inhibitor (SB402451) was sourced from GSK. RNA extraction kits were procured from Qiagen (RNeasy Mini and Micro Kits). anti-RFP antibody (600-401-379) was from Rockland, and anti-collagen type II antibody (MAB8887) was from Merck. Biotinylated goat anti-rabbit (BA-1000) was from Vector. Alexa Fluor 488-conjugated anti-rabbit (ab150065)) and Alexa Fluor 647-conjugated anti-mouse (ab150111) are from Abcam. The reverse transcription kit and polymerase chain reaction (PCR) master mix were obtained from Applied Biosystems.

### Conditioned medium preparation

Porcine articular cartilage was dissected from metacarpophalangeal (MCP) joints from trotters (distal forelimb of 3-6 months old male pigs. Trotters were decontaminated in 2% Virkon. Freshly dissected porcine cartilage was finely chopped for 5 min (cartilage from one trotter in 2 ml) in Dulbecco’s modified Eagle’s medium (DMEM) containing 1% penicillin-streptomycin (pen-strep, Sigma Aldrich) and 1% amphotericin B (ampB, Thermo Fisher Scientific). The resulting medium conditioned by injured cartilage (injury CM) was spun (10 min @3000g). Spun injury CM was stored at -20 ^0^C until further use. For all cell experiments, injury CM was diluted 1:1 with DMEM containing 2% FBS.

### Cell culture

Porcine cartilage was dissected into DMEM supplemented with 10% FBS, 1% pen-strep, and 1% AmpB (referred to as “complete medium”) and digested overnight (18-20h) at 37°C in the presence of 1.0 mg/mL Type II collagenase (Roche). The solution was then passed through a cell strainer (CLS352340, Corning), centrifuged at 1,500 rpm for 5 min at 4°C, and washed twice with serum-free medium. The resulting cells were resuspended in complete medium.

For bulk sequencing and qPCR experiments (Fig. 2A-D): cells were plated at low density (100,000 cells/3.5cm^2^) in complete medium. After 24h, cells were serum-starved for 4h, then pretreated (20mins) with inhibitor (where specified) and stimulated with vehicle control medium (1% FBS), injury CM, injury CM+FGFRi (250 nM), FGFRi (250 nM) alone for different time points. Cells were collected at 4h for RNA isolation.

For IncuCyte experiments (Fig. 3, Fig 5E): porcine chondrocytes were seeded in 48-well plates at low density (30,000 cells/1.9cm^2^). After 24h, cells were serum-starved for 4h, then pretreated (20 min) with inhibitor (where specified) and stimulated control medium (1% FBS), injury CM, injury CM + FGFRi (250 nM), FGFRi (250 nM), Activin A neutralizing antibody (4 µg/mL), or FGF2 neutralizing antibody (4 µg/mL). Cells were monitored for 96h using the IncuCyte® Zoom video microscope (Essen Bioscience). Cell confluence and quantification were measured using the IncuCyte imaging system.

### Gap assay

Gap migration assays were performed in 24-well tissue culture plates. Using sterile forceps, one 2-well silicone insert (Ibidi, Gräfelfing, Germany) was placed per well, ensuring full adhesion to the bottom of the plate. Freshly isolated chondrocytes were resuspended in complete medium and plated into each chamber of the silicone insert (100,000/0.22cm^2^). After 24h, cells were serum-starved for 4h, then pretreated (20 min) with inhibitor (where specified) and stimulated with control medium (1% FBS), injury CM, injury CM + FGFRi (250 nM), or injury CM passed through a heparin-binding column. Cells were visualized at 24h intervals (up to 120 h) using an Olympus CKX41 microscope.

### RNA extraction and real-time quantitative reverse transcription–polymerase chain reaction (qPCR)

Total RNA was extracted from porcine cells and cartilage explants using TRIzol and the Qiagen RNeasy Mini Kit, following the manufacturer’s instructions. Complementary DNA (cDNA) was synthesized from RNA using a high-capacity cDNA Reverse Transcription Kit (Applied Biosystems). Primers for porcine genes were either self-designed or purchased from Taqman and listed in table S3. Real-time PCR was performed with the ViiA 7 system (Applied Biosystems), using SYBR green Master-Mix (Applied Biosystems) or Taqman Master-Mix (Thermal Fisher Scientific), according to the manufacturer’s protocols. Gene expression levels were normalized to 18S rRNA as the internal control, using the ΔΔCt method.

### Chondrogenesis pellet assay

Low-density plated porcine chondrocytes were cultured with injury condition medium for 72h to induce progenitor cells. These progenitor cells were then centrifuged at 240g for 5 min to form cell pellets. The pellets were cultured in 1 mL of defined chondrogenic medium consisting of DMEM with high glucose, 1× L-glutamine, 40 µg/mL L-proline, 25 µg/mL ascorbate-2-phosphate, 100 nM dexamethasone (Sigma), 1× ITS premix (Corning), 100 µg/mL sodium pyruvate (Lonza), with or without 10 ng/mL TGFβ3 (R&D Systems). The medium was refreshed every other day for 10 days. Cell pellets from all groups were collected on day 3 for gene expression analysis and on day 10 for immunohistochemistry staining. Recombinant human activin A or FGF2 were added to the defined chondrogenic medium (including TGFβ) for 3 days and their effects were compared to those of the positive and negative groups for chondrogenic marker gene expression.

### Cartilage injury assays

For cutting injury, cartilage was rapidly dissected from porcine MCP joints, either snap-frozen immediately on dry ice (0h) or cultured as small explants in serum-free DMEM for 4 or 48h before snap-freezing. All cartilage explants were kept at -80 ^0^C until nuclei isolation. For knee articular cartilage impact injury model, full thickness cartilage explants (6mm diameter) were dissected from porcine femoral condyles of 6-9 month old pigs and rested in serum-free medium for 48h. Explants were then subjected to a defined impact energy of 0.2J by a drop tower model as previously described (*50*), or left uninjured. The explants were cultured in DMEM (1% FBS) for a further 2 weeks with either bulk tissue qPCR (0-14 day time points) or snRNA-Seq (14 days only) performed.

### Single Nucleus isolation

For both experiments, cartilage explants were ground by rotational action into small pieces in liquid nitrogen using a cryo-cup grinder, then lysed in a buffer containing 10 mM Tris-HCL (pH 7.4), 10 mM NaCl, 3 mM MgCl2, Tween-20 0.01%, Nonidet P40 substitute 0.1%, Dogitonin 0.01%, and RNAse inhibitor 0.2U). Nuclei were released by Dounce homogenization (grinder A x10, B x10, A x10) (Sigma Aldrich). Upper fraction (containing nuclei) was collected and spun at 500x g for 5 min at 4°C. The resulting pellet was resuspended in wash buffer (PBS 1x, BSA 1%, RNase inhibitor 0.2U), and then filtered through 70 μm and 40 μm SmartStrainer (BAH136800040). The filtered material was centrifuged again, and nuclei pellets were resuspended in wash buffer. Nuclei were stained with DAPI and sorted using FACSAria™ II (BD). The nuclei extraction protocol was repeated on the lower fraction (containing cell debris) of the initial Dounce homogenisation to add to nuclei yield.

### Generation of droplet-based single-nuclei RNA-seq (snRNA-seq) data

Nuclei counts were determined with acridine orange/propidium iodide fluorescence using a LUNA-FX7 cell counter (Logos Biosystems). Approximately 10,000 – 15,000 nuclei per sample were loaded onto the 10X Genomics Chromium Controller (Chip G). Gene expression sequencing libraries were prepared using the 10x Genomics Single Cell 3’ Reagent Kits v3.1 following the manufacturer’s user guide (CG000330). The final libraries were loaded on the NovaSeq6000 sequencing platform (Illumina, v1.5 chemistry, 150bp paired end) at the Oxford Genomics Centre (Centre for Human Genetics, University of Oxford).

### Analysis of snRNA-seq data from injured porcine articular cartilage

10x snRNA-seq data were generated for 10 samples (sample details listed in table S4). Sequencing reads were mapped using Cell Ranger version multi (version 6.0.0) with the Sscrofa11.1 reference transcriptome built using cellranger pipeline. Then CellBender was used to remove ambient UMI counts (*51*). For the analysis we selected nuclei with > 200 genes and < 2% mitochondrial reads (n=31,104 nuclei). To identify doublets in dataset Scrublet Single-Cell Remover of Doublets tool (52) was used with threshold for scrublet score = 0.25 (mean score + 2SD) to predict and filter out problematic multiplets. Data were pre-processed with Scanpy (*53*) (version 1.8.1) (total count normalised and log1p transformed). Highly variable genes (HVG) (n= 4, 331) were identified within the whole data set in at least 1 batch (batches are combination of condition and tissue batch). The effect of total UMI number was regressed out and the data scaled. The data were integrated with python scVI (*54*) (with following parameters n_layers=2, n_latent=30, gene_likelihood="nb"). The integrated data were analysed using pipeline_cluster.py from cellhub (https://github.com/sansomlab/cellhub/). An exact neighbour graph was computed with Scikit-learn (*55*) (as implemented in scVelo) (*56*) using n=30 components, n=20 neighbours and the euclidean distance metric. This neighbour graph was used to compute the UMAP and for Leiden clustering with resolution 0.6. Significant cluster markers were identified using the scanpy “rank_genes_groups” function. Subset of cells identified as chondrocytes (n = 25,075) was used to analyse in depth the relationship between time points. Differential expression analysis (DEA) to compare 4h vs 0h chondrocytes (6 samples listed in table S10) was conducted using the Bioconductor package DESeq2 (version 1.34.0) on pseudobulk counts (57). For the analysis of snRNA-seq data from acute impact injury model, see supplementary methods.

### Bulk RNAseq data analysis

RNA quantity and integrity were assessed with Agilent 5400. Polyadenylation-selected sequencing libraries were prepared by Novogene. Libraries were subjected to 150 paired-end sequencing using Illumina’s NovaSeq X Plus platform. RNAseq analysis was performed using txseq pipelines (https://github.com/sansomlab/txseq). Sequence reads were aligned to the pig genome with HISAT2 (version 2.1.0) (58) using a “genome_trans” index built from the genome assembly Sscrofa11.1 and Ensembl version 111 annotations. Data quality was assessed using pipeline_fastqc.py, The average alignment rate was 82.00%. Mapped reads were counted using featureCounts (Ensembl version 111 annotations, with default parameters) (59). Salmon v0.9.1 was used to calculate transcript per million values (60) using a quasi-index (built with Ensembl version 111 annotations and *k* = 31) and gc bias correction (parameter “--gcBias”).

Differential expression (DE) analysis of CM vs ctrl or CM-FGFRi vs CM samples was performed using Bioconductor package DESeq2 (v1.34.0) (57). Prior to analysis, genes were filtered so as to retain n=17,295 protein-coding genes expressed at sum of counts >0. Gene set enrichment analysis with FGSEA (*61*) was performed using the genes for differential expression by DESeq2 (i.e. after independent filtering, listed in supplementary datafile S3 and S4) using the multilevel procedure with Hallmark pathway annotations (*62*).

#### Murine experiments

All in vivo experiments were carried out under ethical approval in agreement with local policy. Work was conducted under the home office project license (PPL P7E1A3608, University of Oxford) and PDCD943FB for University of Aberdeen. Mice were housed in standard individually ventilated cages under a 12-hour light/12-hour dark cycle. Four- or 10-week-old C57BL/6 and Balb/c mice male mice were obtained from Charles River Laboratories, UK. *Fgf2^−/−^* mice were purchased from Jackson Laboratory. Four- or 10-week-old C57BL/6 and Balb/c mice male mice were obtained from Charles River Laboratories, UK. *Fgf2^−/−^* mice were purchased from Jackson Laboratory. *Col2-CreER* mice (JAX stock no. 6774) (*63*) were on an FVB/N background, while Cre-inducible tdTomato mice (tdTom; JAX stock no. 7914) (*64*) were on a C57BL/6J background. For labelling and tracing of *Col2*-lineage cells, *Col2-CreER* mice were crossed with Cre-inducible *tdTomato* mice. Starting at postnatal day 16, double-transgenic offspring were administered 75 mg/kg of tamoxifen dissolved in corn oil by intraperitoneal injection daily for 5 days to activate CreER. At 10 weeks of age, 15 tamoxifen-induced female mice underwent unilateral focal cartilage injury, performed as previously described (*65*).

Mice undergoing surgery were anesthetized by inhalation of isoflurane (4% induction and 2% maintenance) in *2* litres/min oxygen. All animals received a subcutaneous injection of buprenorphine (Vetergesic; Alstoe Animal Health) peri- and/or post-surgery.

Patella groove focal cartilage defect: An incision was made medially and proximally to the insertion of the patellar tendon and extended proximally to the attachment of the quadriceps muscle. The patella was dislocated laterally and the joint fully flexed to expose the patellar groove. A 25 G needle was placed with its tip just anteriorly to the intercondylar notch in the centre of the groove and dragged proximally across the entire length of the patellar groove repeatedly to create a longitudinal full-thickness cartilage defect. The patellar dislocation was then repositioned and the joint capsule was closed with sutures. The mice were fully mobile within 5 min following withdrawal of Isoflurane. Mice were culled 8 weeks post-surgery and hindlimbs were dissected for histological analysis.

For analysis of *Col2*-lineage contribution following focal cartilage injury, tissue sections from tamoxifen-induced *Col2-CreER* mice that showed evidence of repair, assessed from Safranin O and Fast Green stained sections were selected at set intervals across the joint. Immunohistochemical staining for tdTom was performed to quantify tdTom-expressing *Col2*-lineage cells. Quantification was performed with QuPath v0.3.2 (*66*) in 4-13 sections per knee at set intervals between the start of the full-thickness injury to the region where the growth plate and articular cartilage begin to merge at the peripheral edges. Nine mice showed little or no cellular repair tissue and were excluded from the analysis. Regions of interest (ROIs) were drawn around the repair tissue, the surrounding articular cartilage, subchondral bone marrow, and synovium and periosteum using the wand or brush tool. Cell counting was then performed using the positive cell detector and inaccurate detections deleted or edited manually. The number of cells counted was summed per mouse and the percentage of cells expressing tdTom was calculated.

#### Yap-Tead GFP reporter assay

the generation and validation of C3H10T1/2 Yap-Tead GFP reporter cells, stably transduced with a pGreenFire lentiviral reporter vector containing Tead DNA-binding sequences upstream of an mCMV promoter, is described previously *(32)*. Yap reporter and empty vector control cells were seeded (2,000 cells/cm^2^) in growth media (high-glucose DMEM supplemented with 10% FBS and 4 mM L-glutamine) and allowed to attach for 24 h before overnight serum ‘starvation’ (1% FBS). Cells were then stimulated for 48 h under vehicle-controlled 1% serum conditions as indicated, using recombinant mouse IL-6/IL-6R (20 ng/ml), injury CM (50%, 25% and 10% dilution), and FGFRi (250 nM). GFP fluorescence was detected using a BD Fortessa flow cytometer and analysed using FlowJo v10. To account for background fluorescence, geomean fluorescence intensity was normalized to the empty vector control cells for each treatment condition and shown relative to the vehicle control. All statistical analysis was performed on data before normalization to the vehicle control.

### Immunohistochemistry

Paraformaldehyde-fixed joints from focal cartilage injury operated *Fgf2^-/-^*mice were decalcified in EDTA (2 weeks), embedded in paraffin, sectioned coronally, and stained with Safranin O and hematoxylin and eosin for microscopic inspection and histological scoring. Histological analysis of the knee joints was done as above. For scoring of repair tissue, 3 histological sections were taken from defined anatomical locations. The repair tissue was scored blind, by two independent assessors using a modified Pineda score, where a score of 1 equates to poor repair and 14 equates to complete repair for each section.

Hindlimbs from *Col2-CreER* mice were fixed in 4% paraformaldehyde, decalcified in 10% ethylenediaminetetraacetic acid (EDTA), and paraffin-sectioned at 5 µm thickness. Immunohistochemistry and immunofluorescence staining were performed as previously described (66), using a rabbit anti-RFP primary antibody (600-401-379) and a mouse anti-collagen type II primary antibody (MAB8887), with a biotinylated goat anti-rabbit (BA-1000) or Alexa Fluor 488-conjugated anti-rabbit (ab150065) and Alexa Fluor 647-conjugated anti-mouse (ab150111) secondary antibodies. Images were acquired on a Zeiss Axioscan Z1 or Zeiss LSM710 confocal microscope with ZEN software.

#### Analysis of Protein Changes analysed by SomaScan Following Joint Distraction

The Knee Joint distraction cohort (ethics number: 15-160/D; NL51539.041.15) comprised n = 20 participants identified by the orthopaedic surgeon from a population of symptomatic knee OA patients attending clinic for consideration of KJD as part of their usual clinical care (Utrecht, Netherlands) (*34*). Key inclusion criteria included: aged <65 years, knee <u>OA</u> fulfilling ACR clinical criteria and Kellgren and Lawrence (KL) grade≥2, details are listed in table S5. As part of the KJD cohort, SF was collected from the index knee at the baseline visit (prior to the distraction frame being fitted), at 3-4 weeks, and at 6-7 weeks (immediately after the distraction frame was removed under anaesthesia). For this analysis, only participants with paired SF samples at baseline and 6 weeks were included. A single pair was excluded due to issues with protein reads, leaving 16 pairs (n = 32 samples).

All raw protein data (received from SomaLogic) underwent our in-house quality control procedure (*67*), which briefly, included standardisation, technical confounder correction and filtering. As part of the standardisation, batch-correction for spin-status using the *ComBat* function in R (version 0.0.4) (*67,68*) was also applied. We generated volcano plots to visualize the proteins associated with the paired t-test analyses, labelling the top 15 positive and negatively regulated based on adjust p value. A limited number of proteins had multiple SOMAmers on the platform so for these cases, only the most significant SOMAmer, based on the ranked adjusted p-value, was labelled on the volcano plots. To investigate change in protein expression resulting from joint distraction, we employed paired t-tests where each SOMAmer was modelled individually. Prior to fitting the models, protein expression values were transformed using natural logarithms. We then standardized these values on a per-protein basis within the Discovery dataset by subtracting the mean log protein abundance and dividing by the standard deviation. The coefficients derived from the paired t-tests reflect the mean change in log protein abundance for the given protein upon joint distraction. We adjusted p-values for multiple comparison testing using the Benjamini-Hochberg correction method. Pathway Enrichment Analysis of SomaScan data see supplementary methods.

#### Statistical analysis

Data are expressed as the means ± SEM or minimum to maximum with mean for replicate experiments. Analysis was performed using Prism 10 software (GraphPad). Student’s *t* tests were used to establish statistical significance between two groups. One-way analysis of variance (ANOVA) was used to compare multiple groups. Two-way ANOVA with Bonferroni’s posttest was performed for multiple comparisons. *P* values less than 0.05 were considered statistically significant unless otherwise stated. Other post hoc testing was done as specified in figure legends.

## Supplementary Materials

Materials and Methods

Fig. S1 to S4

Tables S1-S5

Data file S1-S7

## Acknowledgement

The STEpUP OA Consortium author block includes: University of Nottingham: Ana M. Valdes, David A. Walsh, Michael Doherty, Vasileios Georgopoulos; Lund University: Staffan Larsson, L. Stefan Lohmander, André Struglics; University of Cambridge: Brian D.M. Tom, Laura Bondi; University of Toronto: Mohit Kapoor, Rajiv Gandhi, Anthony Perruccio, Y. Raja Rampersaud, Kim Perry; University of Manchester: Tim Hardingham, David Felson; University of Oxford: Tonia L. Vincent, Thomas A. Perry, Luke Jostins-Dean, Yun Deng, Vicky Batchelor, Jennifer Mackay-Alderson, Gretchen Brewer, Rose M. Maciewicz, Brian Marsden, Nigel K. Arden, Philippa Hulley, Andrew Price, Stefan Kluzek, Megan Goff, Vinod Kumar, James Tey; Imperial College London: Fiona E. Watt, Andrew Williams, Artemis Papadaki; University College Maastricht: Tim J. Welting, Pieter Emans, Tim Boymans, Liesbeth Jutten, Marjolein Caron, Guus van den Akker; University of Western Ontario: C. Thomas Appleton, Trevor B. Birmingham, J. Daniel Klapak; Biosplice: Sarah Kennedy, Jeymi Tambiah; Fidia: Devis Galesso, Nicola NK; SomaLogic: Joe Gogain, Darryl Perry, Anna Mitchel, Ela Zepko; Novartis: Sophie Brachat, Joanna Mitchelmore, Juerg Gasser, Lori Jennings; UCB: Waqar Ali. We thank the orthopedic surgeon Dr R Custers, Department of orthopaedics, UMC Utrecht, Netherlands, for synovial fluid collection and SJD surgeries. We thank colleagues from the Kennedy histology laboratory, I. Parisi and B. Stott, for their support.

## Funding

German Academic Exchange Service DAAD (JR)

Dutch Arthritis Society LLP-9 (SCM)

ZonMW-NWO talent program VENI (MJ)

UKRI Future Leaders Fellowship MR/S016538; MR/S016538/2; MR/Y003470/1 (FEW)

Centre for OA Pathogenesis Versus Arthritis Grants 20205 and 21621 (TLV)

Versus Arthritis project grant 20783 (TLV)

Reumafonds grant ISP14-3-301/16-1-404 (SCM)

Tissue Engineering and Regenerative Therapies Centre Versus Arthritis grant 21156 (CDB, AJR)

Versus Arthritis project grant 20775 (CDB, AJR)

## Author contributions

Conceptualization: LZ, HM, SNS, JR, CDB, AJR, TLV

Methodology: LZ, HM, TAP, YI, JZM, STEpUP OA Consortium, FEW, YI, JZM, MA

Investigation: LZ, HM, TAP, AA, RAS, JJM, KK, SMC, FC, LAK, SK, SEE, FDA, SM, MJ, JB, CC, JR

Visualization: LZ, LAD, TAP

Funding acquisition: TLV, CDB, AJR, FEW, SCM, JR

Writing – original draft: LZ, TLV

Writing – review & editing: LZ, HM, TAP, AA, RAS, JJM, KK, SMC, LAK, SK, SEE, FEW, FDA, SM, CC, JR, CDB, AJR, TLV

## Data and materials availability

All data are available in the main text, supplementary materials, or the listed GEO files: GSE291306 (snRNAseq drop tower model), GSE291191(bulkRNAseq) and GSE291579 (snRNAseq cutting injury model).

