## supplemental method for "Mechano-activated chondroprogenitors (MACs) drive intrinsic cartilage repair; a process that is arrested in osteoarthritis"

**This PDF file includes:**

Materials and Methods

Figs. S1 to S4

Tables S1-S5

**Other Supplementary Materials for this manuscript include the following:**

Data file S1-S7

**Materials and Methods**

**Analysis of snRNA-seq data from acute impact injury model**

The 10x snRNA-seq data were generated for 6 samples (3 ctrl and 3 trauma, sample details listed in table S10). Sequencing reads were mapped using Cell Ranger version multi (version 7.1.0) with the Sscrofa11.1 reference transcriptome built using cellhub pipeline (https://github.com/sansomlab/cellhub). For the analysis we selected nuclei with > 200 genes and < 2% mitochondrial reads (n=24,782 nuclei). Doublets were identified using the method above. Data were pre-processed with Scanpy (version 1.9.8) (total count normalised and log1p transformed). Highly variable genes (HVG) were identified within the data set (n=2000). The effect of total UMI number was regressed out and the data scaled. The data were integrated and clustered as described for the injured porcine articular cartilage. The neighbour graph was used to compute the UMAP and for Leiden clustering with resolution 0.6. Significant cluster markers were identified using the scanpy “rank_genes_groups” function. Subset of cells identified as chondrocytes (n = 24,022) was used to analyse in depth the relationship between time points. Differential expression analysis (DEA) to compare injured vs rested chondrocytes was conducted using the Bioconductor package DESeq2 on pseudobulk counts.

**Pathway Enrichment Analysis of SomaScan data**

To assess the enrichment of the associated proteins within biological pathways, we utilized gene sets from The Molecular Signatures Database (MSigDB) focusing on Hallmark, and Reactome (69). Proteins were mapped to the respective gene sets using 'EntrezGeneSymbol', 'Target', or 'EntrezGeneID' variables provided by SomaLogic. We performed protein set enrichment testing using the *fgsea* package in R (version 1.28.0) to identify pathways with genes significantly enriched in association with knee joint distraction. All proteins included in the respective analyses were ranked based on a computed 'rank metric', defined as the product of -log(p-values) and the sign of the coefficients. Finally, we utilized the *ggplot2* package in R (version 3.5.0) to create bubble plots and line plots for visualizing the results.

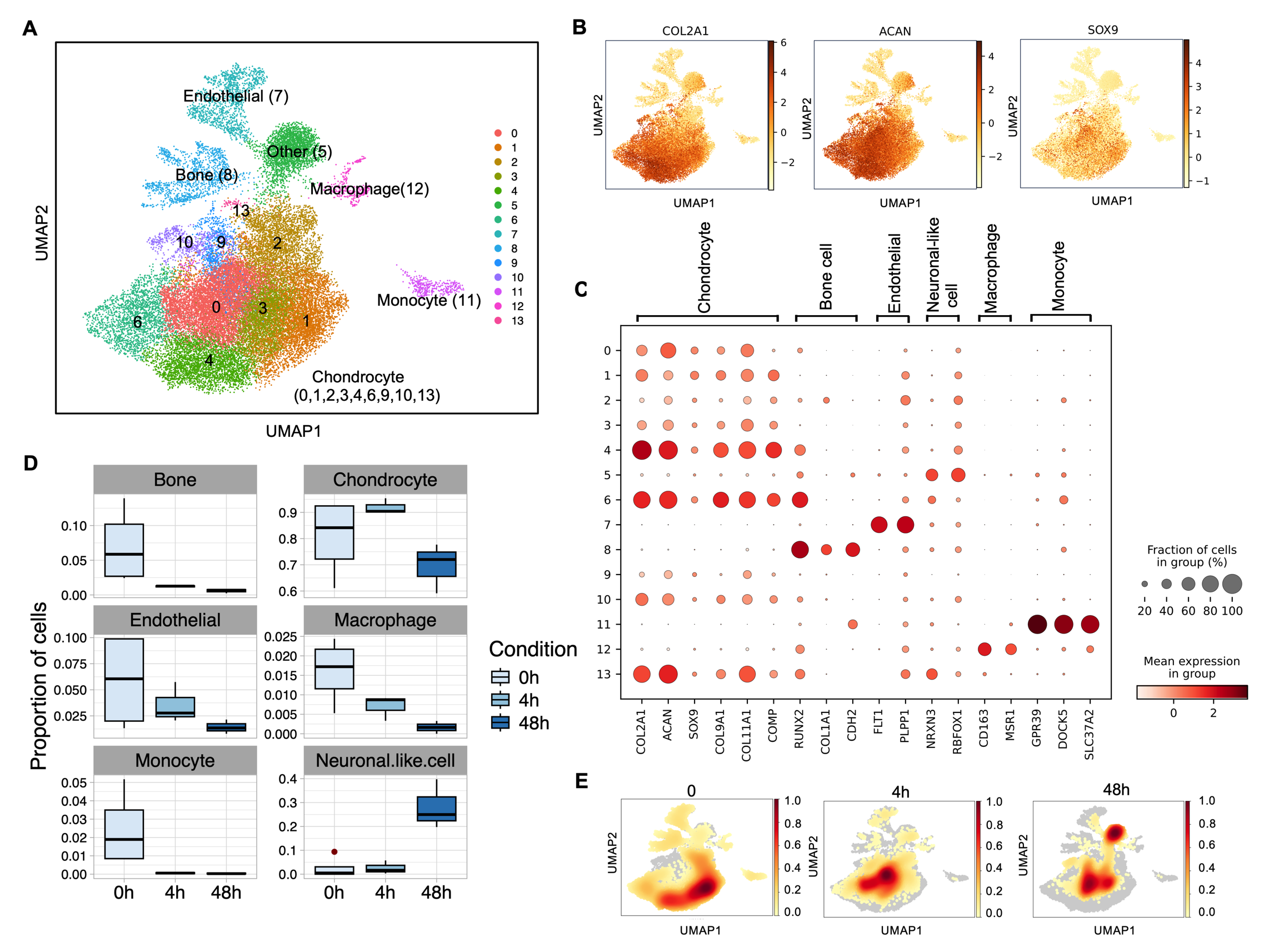

**Fig. S1. Single-nuclei RNA-seq of porcine articular cartilage upon cutting injury.** **A,** Uniform manifold approximation and projection (UMAP) plot of all cells extracted from cartilage explants (*n* = 31,104 nuclei; 0h, n=4; 4h, n=3; and 48h, n=3), coloured by cluster and labelled by cell type. **B,** UMAP showing the expression of chondrocyte marker genes *COL2A1, ACAN*, and *SOX9*. *COL2A1* – Collagen Type II Alpha 1 Chain; *ACAN*- Aggrecan, *SOX9*- SRY-Box Transcription Factor 9. **C**, Dot plot of the top marker genes (X-axis) for each cell cluster (Y-axis), with the relative fraction of cells expressing gene and level of expression shown. **D**, The boxplots show the proportion of cells in each cluster calculated as percentage of the total cells at each condition. **E**, Density plot of cells across different time points superimposed on the UMAP representation.

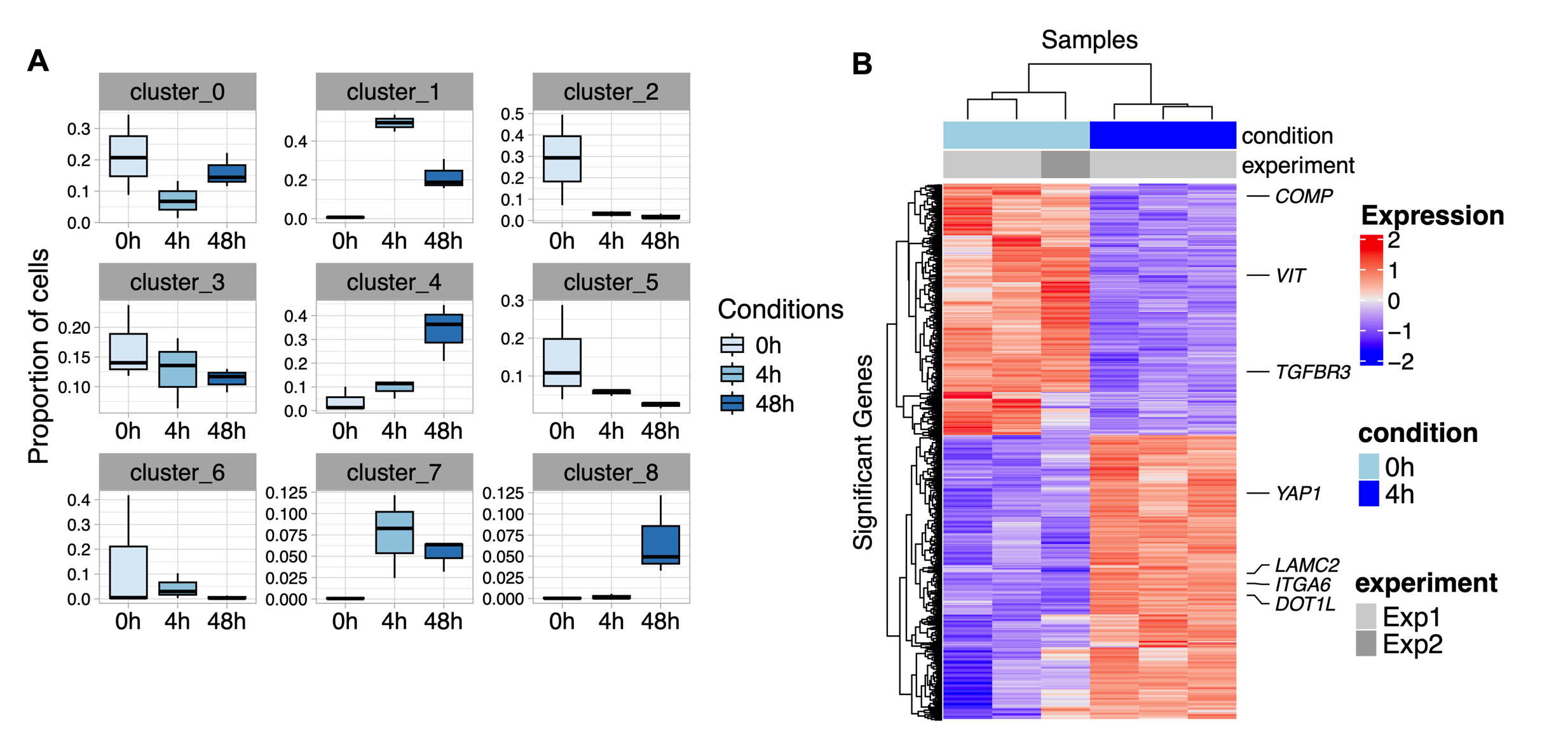

**Fig S2. Single-nuclei RNA-seq of porcine chondrocytes after injury. A**, Proportion of cells in each cluster calculated as percentage of the total cells at each condition (0h, n=3; 4h, n=3, sample details listed in table S10). **B**, Heatmap of row scaled pseudobulk counts showing a set of putative progenitor cell cluster marker genes (*ITGA6, LAMC2, YAP1,* and *DOT1L*) as well as the negative marker genes for Cluster 1 (*TGFBR3, VIT* and *COMP*). *TGFBR3* – TGFβ receptor 3; *VIT* – vitronectin; *COMP* – cartilage oligomeric protein; *ITGA6* – integrin alpha-6; *YAP1* – yes-associated protein-1; *DOT1L* – DOT1-like histone lysine methyltransferase (Deseq2 analysis, BH adjust p < 0.05, |fold change| > 1.5). For other gene identities see Supplementary Data File 1.

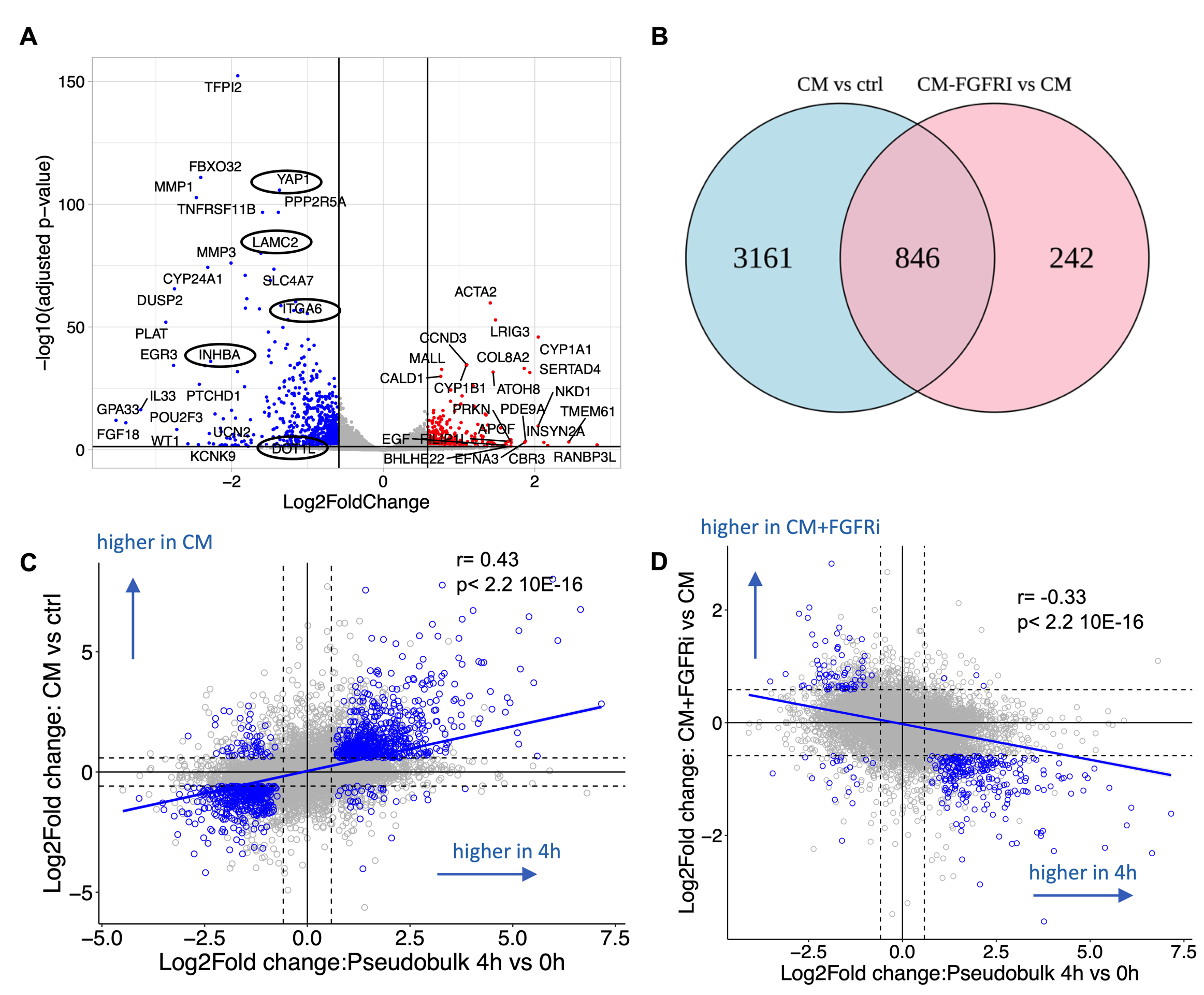

**Fig S3. FGFR-dependent contribution to injury CM gene regulation.** **A**, Enhanced volcano plot showing differentially expressed chondrocyte genes (DEG) following stimulation with injury CM (CM) with FGFR inhibition (FGFRi) compared with injury CM alone, padj < 0.05 and |Foldchange|> 1.5. Red dots indicate upregulated genes when FGFRi is included; blue dots indicate genes downregulated by FGFRi. **B**,Venn diagram showing overlap of genes which were regulated by injury CM with those that were FGFR-dependent. Scatter plots showing the relationship between genes identified in snRNAseq pseudobulk DEGs after cartilage injury (4h vs 0h) with **C**, DEGs regulated by chondrocytes stimulated with injury CM (compared with control) or **D**, DEGs regulated by the addition of FGFRi to injury CM-stimulated chondrocytes. Blue dots represented the genes with |Foldchange|> 1.5 and padj<0.05.

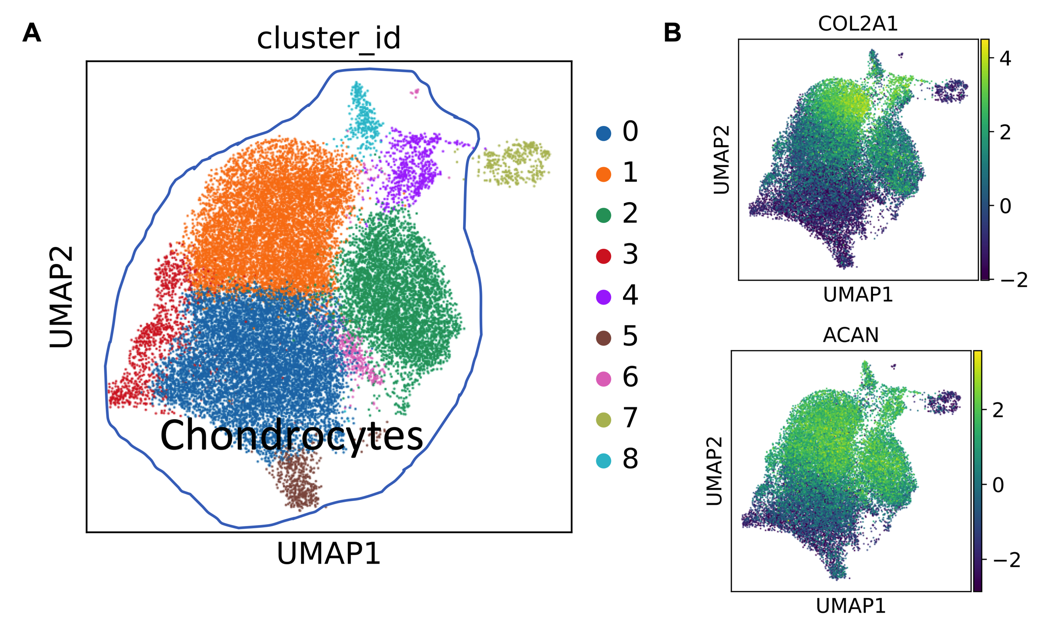

**Fig S4. Single-nuclear RNA-seq of porcine articular cartilage upon drop tower impact injury.** **A**, Uniform manifold approximation and projection (UMAP) plot of all cells isolated from knee cartilage explants (*n* = 24,782 nuclei; ctrl, n=3; trauma, n=3), coloured by cluster and chondrocytes clusters were labelled. **B**, UMAP showing the expression of chondrocyte marker genes *ACAN*, and *COL2A1*.

**Table S1. Significant regulated pathways by gene enrichment analysis (Hallmark) of injury CM vs ctrl. FGSEA analysis, BH adj. p < 0.05.**

| **pathway** | **pval** | **padj** | | **log2err** | | | **ES** | **NES** | **Size** |
| --- | --- | --- | --- | --- | --- | --- | --- | --- | --- |
| Hallmark_Tnfa_Signaling_Via_Nfkb | 5.60E-32 | | 2.80E-30 | | 1.468 | 0.774 | | 2.509 | 181 |
| Hallmark_Inflammatory_Response | 2.57E-18 | | 6.43E-17 | | 1.105 | 0.720 | | 2.289 | 143 |
| Hallmark_Epithelial_Mesenchymal_Transition | 3.06E-14 | | 5.10E-13 | | 0.965 | 0.645 | | 2.090 | 185 |
| Hallmark_Il2_Stat5_Signaling | 6.73E-14 | | 8.41E-13 | | 0.955 | 0.659 | | 2.120 | 167 |
| Hallmark_Myc_Targets_V1 | 1.07E-13 | | 1.07E-12 | | 0.955 | 0.645 | | 2.087 | 177 |
| Hallmark_Hypoxia | 7.58E-13 | | 6.32E-12 | | 0.921 | 0.639 | | 2.072 | 181 |
| Hallmark_Myc_Targets_V2 | 4.26E-12 | | 3.04E-11 | | 0.887 | 0.802 | | 2.305 | 52 |
| Hallmark_Kras_Signaling_Up | 1.11E-10 | | 6.92E-10 | | 0.839 | 0.624 | | 2.004 | 165 |
| Hallmark_Il6_Jak_Stat3_Signaling | 5.37E-10 | | 2.98E-09 | | 0.801 | 0.750 | | 2.186 | 63 |
| Hallmark_Allograft_Rejection | 3.03E-09 | | 1.52E-08 | | 0.775 | 0.626 | | 1.986 | 136 |
| Hallmark_Estrogen_Response_Early | 1.49E-07 | | 6.76E-07 | | 0.690 | 0.569 | | 1.838 | 175 |
| Hallmark_Coagulation | 3.67E-06 | | 1.33E-05 | | 0.627 | 0.594 | | 1.847 | 105 |
| Hallmark_Unfolded_Protein_Response | 3.73E-06 | | 1.33E-05 | | 0.627 | 0.600 | | 1.871 | 106 |
| Hallmark_Uv_Response_Dn | 3.41E-06 | | 1.33E-05 | | 0.627 | 0.562 | | 1.782 | 135 |
| Hallmark_Complement | 5.26E-06 | | 1.75E-05 | | 0.611 | 0.560 | | 1.783 | 152 |
| Hallmark_Apoptosis | 1.35E-05 | | 4.21E-05 | | 0.593 | 0.545 | | 1.735 | 143 |
| Hallmark_Interferon_Gamma_Response | 2.07E-05 | | 6.09E-05 | | 0.576 | 0.532 | | 1.696 | 158 |
| Hallmark_Mtorc1_Signaling | 2.94E-05 | | 8.16E-05 | | 0.576 | 0.514 | | 1.665 | 185 |
| Hallmark_Tgf_Beta_Signaling | 3.62E-05 | | 9.52E-05 | | 0.557 | 0.666 | | 1.908 | 51 |
| Hallmark_P53_Pathway | 5.23E-05 | | 0.00013067 | | 0.557 | 0.510 | | 1.650 | 179 |
| Hallmark_Estrogen_Response_Late | 5.82E-05 | | 0.00013855 | | 0.557 | 0.521 | | 1.674 | 167 |
| Hallmark_Uv_Response_Up | 0.00034957 | | 0.00079447 | | 0.498 | 0.510 | | 1.617 | 135 |
| Hallmark_Apical_Surface | 0.0003816 | | 0.00082957 | | 0.498 | 0.681 | | 1.804 | 35 |
| Hallmark_Angiogenesis | 0.00046672 | | 0.00097232 | | 0.498 | 0.688 | | 1.799 | 32 |
| Hallmark_Apical_Junction | 0.00052321 | | 0.00104641 | | 0.477 | 0.483 | | 1.556 | 171 |
| Hallmark_Glycolysis | 0.00393619 | | 0.0075696 | | 0.407 | 0.448 | | 1.443 | 171 |
| Hallmark_Myogenesis | 0.00638818 | | 0.01182995 | | 0.407 | 0.451 | | 1.438 | 158 |
| Hallmark_Androgen_Response | 0.00741011 | | 0.01323234 | | 0.407 | 0.512 | | 1.559 | 88 |
| Hallmark_Reactive_Oxygen_Species_Pathway | 0.01014138 | | 0.01748513 | | 0.381 | 0.582 | | 1.604 | 42 |
| Hallmark_G2m_Checkpoint | 0.02285514 | | 0.0380919 | | 0.352 | 0.415 | | 1.346 | 186 |

**Table S2. Significant regulated pathways by gene enrichment analysis (Hallmark) of injury CM with FGFRi vs injury CM. FGSEA analysis, BH adj. p < 0.05.**

| **pathway** | **pval** | **padj** | **log2err** | **ES** | **NES** | **size** |
| --- | --- | --- | --- | --- | --- | --- |
| Hallmark_Tnfa_Signaling_Via_Nfkb | 1.66E-23 | 8.28E-22 | 1.255 | -0.684 | -2.503 | 174 |
| Hallmark_E2f_Targets | 1.24E-22 | 3.11E-21 | 1.230 | 0.579 | 2.904 | 185 |
| Hallmark_Inflammatory_Response | 4.02E-14 | 6.69E-13 | 0.965 | -0.650 | -2.324 | 124 |
| Hallmark_Il2_Stat5_Signaling | 4.66E-12 | 5.82E-11 | 0.887 | -0.609 | -2.206 | 152 |
| Hallmark_Epithelial_Mesenchymal_Transition | 2.04E-09 | 2.04E-08 | 0.775 | -0.544 | -1.997 | 178 |
| Hallmark_Kras_Signaling_Up | 6.72E-09 | 5.34E-08 | 0.761 | -0.558 | -2.017 | 145 |
| Hallmark_Uv_Response_Dn | 7.48E-09 | 5.34E-08 | 0.748 | -0.577 | -2.072 | 131 |
| Hallmark_G2m_Checkpoint | 1.47E-07 | 9.17E-07 | 0.690 | 0.391 | 1.961 | 185 |
| Hallmark_Hypoxia | 1.94E-07 | 9.72E-07 | 0.690 | -0.521 | -1.904 | 167 |
| Hallmark_Oxidative_Phosphorylation | 1.93E-07 | 9.72E-07 | 0.690 | 0.382 | 1.895 | 177 |
| Hallmark_Il6_Jak_Stat3_Signaling | 3.19E-07 | 1.45E-06 | 0.675 | -0.663 | -2.122 | 55 |
| Hallmark_Coagulation | 7.72E-06 | 3.22E-05 | 0.593 | -0.550 | -1.909 | 97 |
| Hallmark_Androgen_Response | 5.80E-05 | 0.000209 | 0.557 | -0.536 | -1.825 | 83 |
| Hallmark_Estrogen_Response_Early | 5.86E-05 | 0.000209 | 0.557 | -0.469 | -1.708 | 159 |
| Hallmark_Complement | 0.000122 | 0.000405 | 0.538 | -0.471 | -1.697 | 138 |
| Hallmark_Apical_Junction | 0.000159 | 0.000498 | 0.519 | -0.471 | -1.708 | 149 |
| Hallmark_Interferon_Gamma_Response | 0.000267 | 0.000786 | 0.498 | -0.468 | -1.686 | 134 |
| Hallmark_Allograft_Rejection | 0.000323 | 0.000849 | 0.498 | -0.471 | -1.671 | 116 |
| Hallmark_Apoptosis | 0.000316 | 0.000849 | 0.498 | -0.468 | -1.683 | 135 |
| Hallmark_Cholesterol_Homeostasis | 0.000546 | 0.001365 | 0.477 | -0.532 | -1.751 | 69 |
| Hallmark_P53_Pathway | 0.000881 | 0.002097 | 0.477 | -0.422 | -1.546 | 174 |
| Hallmark_Estrogen_Response_Late | 0.000947 | 0.002153 | 0.477 | -0.442 | -1.603 | 151 |
| Hallmark_Mtorc1_Signaling | 0.002237 | 0.004863 | 0.432 | -0.408 | -1.503 | 182 |
| Hallmark_Myc_Targets_V2 | 0.003377 | 0.007036 | 0.432 | -0.536 | -1.692 | 52 |
| Hallmark_Notch_Signaling | 0.003892 | 0.007783 | 0.432 | -0.606 | -1.724 | 28 |
| Hallmark_Pi3k_Akt_Mtor_Signaling | 0.005736 | 0.01103 | 0.407 | -0.455 | -1.556 | 87 |
| Hallmark_Glycolysis | 0.007277 | 0.013475 | 0.407 | -0.397 | -1.448 | 163 |
| Hallmark_Myogenesis | 0.008106 | 0.014476 | 0.381 | -0.416 | -1.497 | 135 |
| Hallmark_Apical_Surface | 0.010901 | 0.018794 | 0.381 | -0.564 | -1.649 | 31 |
| Hallmark_Angiogenesis | 0.019101 | 0.030808 | 0.352 | -0.546 | -1.594 | 31 |
| Hallmark_Dna_Repair | 0.018553 | 0.030808 | 0.352 | 0.292 | 1.393 | 130 |
| Hallmark_Hedgehog_Signaling | 0.028444 | 0.044444 | 0.352 | -0.558 | -1.554 | 26 |

**Table S3. Primers and Taqman probes used for Real-Time PCR.**

| **Gene ID** | **Probe ID/ primer sequence** |
| --- | --- |
| *18s* | Hs99999901_s1 |
| *ACAN* | Ss03374823_m1 |
| *SOX9* | Ss03392406_m1 |
| *COL1A1* | Ss03373340_m1 |
| *COL2A1* | Ss03373344_g1 |
| *YAP1* | Ss06912139_m1 |
| *ITGA6* | F-5'-AGAGCCAATCACAGCGGAG-3'; R-5'-GCCAAATGAAGAAGCCAGCC-3' |
| *LAMC2* | F-5'-CTGGGACCTTTTGGCACCT-3'; R-5'-CTTCCTCTGTCTCGGGCATC-3' |
| *18s* | F-5'-TGCAGAATCCTCGCCAATACA-3'; R-5'-AGTCGCTCCAAGTCTTCACG-3' |

**Table S4. Sample details for the two snRNAseq data sets.**

* Sample was excluded from the Pseudobulk differential expression analysis (Supplementary Fig. S2b) and formal composition analysis (Supplementary Fig. S2a, S3a, S3b) because sample 0h_1 and 0h_2 were extracted from the same trotter in the tissue batch 1.

| **sampleID** | **joint** | **tissue batch** | **condition** | **data_set** | **number of nuclei** |
| --- | --- | --- | --- | --- | --- |
| 0h_1 | trotter_1 | 1 | 0h | cut_injury | 3990 |
| 0h_2* | trotter_1 | 1 | 0h | cut_injury | 3855 |
| 0h_3 | trotter_2 | 2 | 0h | cut_injury | 3210 |
| 0h_4 | trotter_3 | 2 | 0h | cut_injury | 3812 |
| 4h_1 | trotter_2 | 2 | 4h | cut_injury | 1949 |
| 4h_2 | trotter_3 | 2 | 4h | cut_injury | 2703 |
| 4h_3 | trotter_4 | 2 | 4h | cut_injury | 3392 |
| 48h_1 | trotter_2 | 2 | 48h | cut_injury | 1270 |
| 48h_2 | trotter_3 | 2 | 48h | cut_injury | 3354 |
| 48h_3 | trotter_4 | 2 | 48h | cut_injury | 3074 |
| ctrl_1 | knee_1 | 3 | ctrl | drop tower | 6426 |
| ctrl_2 | knee_2 | 3 | ctrl | drop tower | 2782 |
| ctrl_3 | knee_3 | 3 | ctrl | drop tower | 6181 |
| trauma_1 | knee_1 | 3 | trauma | drop tower | 3948 |
| trauma_2 | knee_2 | 3 | trauma | drop tower | 2898 |
| trauma_3 | knee_3 | 3 | trauma | drop tower | 2547 |

**Table S5. Baseline demographic and clinical characteristics for N = 16 joint distraction participants (with paired synovial fluid sampling at baseline and 6-weeks follow-up).**

| **Feature** | **Feature Description** | **Median (IQR) or n (%)** |
| --- | --- | --- |
| Age | Participant age at the time of sampling (year) | 54.5 (7.25) |
| Sex | Biological sex | Female = 6 (37.5%) |
|  |  | Male = 10 (62.5%) |
| BMI | Participant body mass index at the time of sampling | 29.18 (5.55) |
| WOMAC Pain Score | Scale of 0-100, where 100 is the worst possible knee pain | 57.5 (20) |
| Advanced Radiographic Status | Binary indicator for the presence of advanced stage radiographic knee OA (KL grades 3-4) | Non-advanced: n = 1 (6.25%) |
|  |  | Advanced: n = 15 (93.75%) |
| Ordinal KL Grade | Kellgren Lawrence grade (0-4) (worst affected compartment) | 0: 0 (0%) |
|  |  | 1: 0 (0%) |
|  |  | 2: 1 (6.25%) |
|  |  | 3: 9 (56.25%) |
|  |  | 4: 6 (37.50%) |
| Spin Status | Binary indicator for whether SF sample had been centrifuged prior to supernatant storage | Spun: n = 32 (100%) |

**Data file S1.** Marker genes for snRNAseq data in Figure 1.

**Data file S2.** Differentially expressed genes of the pseudobulk counts of sRNAseq data at 4h vs 0h.

**Data file S3.** Significantly differentially expressed genes following stimulation with injury CM compared with vehicle control.

**Data file S4.** Significantly differentially expressed genes following stimulation with injury CM with FGFRi compared with injury CM alone.

**Data file movie 5.** Live imaging assessed by IncuCyte imaging (at 2h intervals) over 96h. Movie showing change in cell morphology and proliferation for each condition.

**Data file S6.** Significantly expressed genes of the pseudobulk counts of sRNAseq data 2 weeks post injury trauma vs ctrl.

**Data file S7.** Proteins regulated in Joint Distraction synovial fluid.
